# Resolution of ring chromosomes, Robertsonian translocations, and complex structural variants from long-read sequencing and telomere-to-telomere assembly

**DOI:** 10.1101/2023.09.07.555775

**Authors:** Yulia Mostovoy, Philip M. Boone, Yongqing Huang, Kiran V. Garimella, Kar-Tong Tan, Bianca E. Russell, Monica Salani, Benjamin Curall, Diane Lucente, Tera Bowers, Tim DeSmet, Stacey Gabriel, Cynthia C. Morton, Matthew Meyerson, James Gusella, Fabiola Quintero-Rivera, Harrison Brand, Michael E. Talkowski

**Affiliations:** Center for Genomic Medicine, Massachusetts General Hospital, Boston, MA; Program in Medical and Population Genetics, Broad Institute of MIT and Harvard, Cambridge, MA; Department of Neurology, Massachusetts General Hospital and Harvard Medical School, Boston, MA; Division of Genetics and Genomics, Boston Children’s Hospital, Boston, MA; Data Sciences Platform, Broad Institute of MIT and Harvard, Cambridge, MA; Department of Medical Oncology, Dana-Farber Cancer Institute, Boston, MA 02215, USA; Cancer Program, Broad Institute of MIT and Harvard, Cambridge, MA 02142, USA; Division of Genetics, Department of Pediatrics, University of California Los Angeles, Los Angeles, CA; Genomics Platform, Broad Institute of MIT and Harvard, Cambridge, MA; Division of Obstetrics and Gynecology, Brigham and Women’s Hospital, Boston, MA; Department of Genetics, Harvard Medical School, Boston, MA; Departments of Pathology, Laboratory Medicine, and Pediatrics, Division of Genetic and Genomic Medicine, University of California Irvine, Irvine, CA; Pediatric Surgery Research Laboratory, Department of Pediatrics, Boston, MA; Stanley Center for Psychiatric Research, Broad Institute of MIT and Harvard, Cambridge, MA

**Keywords:** Telomere-to-telomere, ring chromosome, Robertsonian translocation, acrocentric p-arm, long-read sequencing, inversion

## Abstract

The capacity to resolve structural variants (SVs) at sequence resolution in highly repetitive genomic regions has long been intractable. Consequently, the properties, origins, and functional effects of multiple classes of complex rearrangement are unknown. To resolve these challenges, we leveraged recent technical milestones: 1) Oxford-Nanopore (ONT) sequencing; 2) the gapless Telomere-to-Telomere (T2T) genome assembly; and 3) a novel tool to discover large-scale rearrangements from long-reads. We applied these technologies across 13 patients with ring chromosomes, Robertsonian translocations, and complex balanced SVs that were unresolved by short-read sequencing. We resolved 10 of 13 events, including ring chromosomes, the complex SVs, and a Robertsonian translocation. Multiple breakpoints were localized to highly repetitive regions inaccessible to short-read alignment, such as acrocentric p-arms, ribosomal DNA arrays, and telomeric repeats, and involved complex structures such as a deletion-inversion and interchromosomal dispersed duplications. We also leveraged ONT native methylation detection to discover phased differential methylation in a gene promoter proximal to a ring fusion site, suggesting a long-range positional effect with heterochromatin spreading. Breakpoint sequences were consistent with common mechanisms of SV formation, including microhomology-mediated mechanisms, non-homologous end-joining, and non-allelic homologous recombination. These methods provide some of the first glimpses into the sequence resolution of ring chromosomes and Robertsonian translocations and illuminate the structural diversity of chromosomal rearrangements with implications for molecular diagnosis and genome biology.

**Highlights:**

- Cryptic rearrangements revealed in repetitive and previously inaccessible regions
- Aligning long-reads to telomere-to-telomere assembly enabled breakpoint discovery
- First sequence resolution of a Robertsonian translocation breakpoint
- Haplotype-specific methylation changes associated with ring formation

**Graphical abstract:** 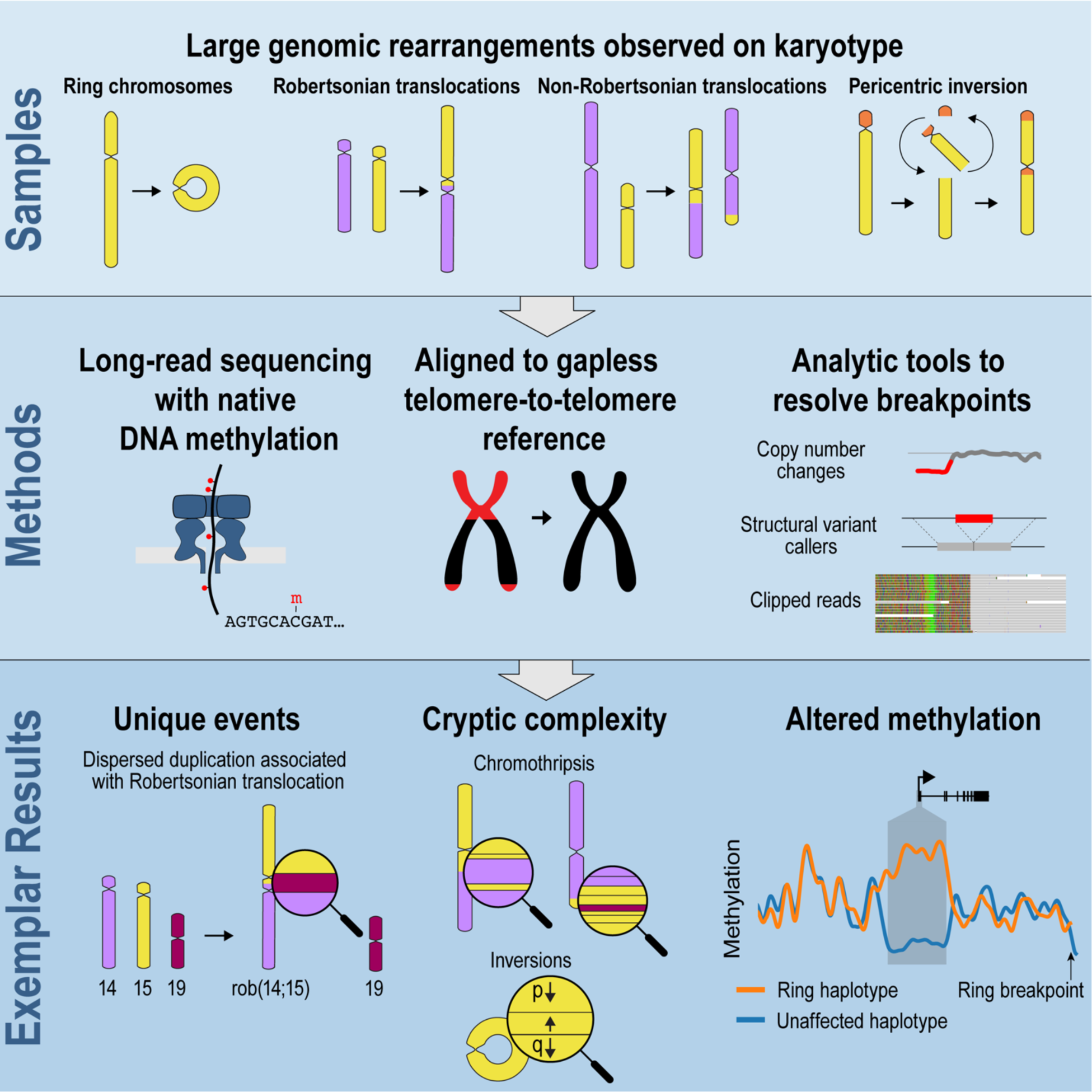

## Introduction

In the emerging field of cytogenomics, an array of molecular and computational methods have been developed to capture chromosomal rearrangements at near sequence resolution, including high-density microarrays^1^, targeted capture, and short-read sequencing (srGS), as well as unselected and genome-wide long-insert^2–5^ and short-insert srGS^6,7^; however, these methods have not provided a panacea to delineate all classes of rearrangement breakpoints. Advances in srGS analytic methods have enabled reliable detection of coding and noncoding single nucleotide variants (SNVs), indels, and many classes of canonical and complex structural variants (SVs)^8^. Despite these advances, highly repetitive genomic regions remain largely recalcitrant to breakpoint identification from srGS^9^, and the most elusive variants have been those with breakpoints that are enriched in long, highly repetitive regions such as acrocentric p-arms, centromeres, and telomeres. These include several classes of chromosomal rearrangements that are commonly observed by karyotyping, including Robertsonian translocations, ring chromosomes, and SVs with breakpoints in classically impenetrable genomic sequences such as repeat arrays and centromeres. Indeed, as clinical genetic testing advances toward nucleotide resolution from exome or srGS as a first-tier diagnostic screen for many rare diseases and developmental disorders, discovery of these large and often deleterious classes of SVs remains a major void in genomic medicine.

Ring chromosomes are circular structures that present in one of three forms: complete ring with rearrangement breakpoints in telomeres/subtelomeres and negligible loss of chromosomal material; ring with terminal chromosomal deletion(s) and/or additional rearrangements(s); or small supernumerary ring chromosome^10,11^. The phenotypic consequence depends on the chromosome involved and the genes that are potentially altered. For example, r(20) is associated with epilepsy^12^, and r(17) with terminal 17p deletion causes Miller-Dieker syndrome from loss of *LIS1*, *YWHAE*, and potentially other genes (https://omim.org/entry/247200). More broadly, a “ring chromosome syndrome” has been observed across rings of various chromosomes, involving poor growth and developmental delay^13^ potentially related to mitotic errors and secondary chromosome rearrangements from post-replication catenation of rings^11^. Healthy carriers with incidentally identified ring chromosomes exist^14^, and these individuals are at risk for infertility owing to meiotic errors^15^.

Robertsonian translocations involve a joining of acrocentric chromosomes in their p-arms or centromeres^16^. Acrocentric p-arms are megabases in length, containing variable numbers of alpha-satellites, 18S/5.8S/28S ribosomal RNA (rRNA) genes coinciding with nucleolar organizing regions (NORs), and beta-satellites among other tandem repeats^11,17,18^. Rearrangements can include p-terminal deletions ranging up to the entire p-arm of each chromosome. Contrasting the clinical impact of ring chromosomes, healthy carriers that are unaware of the Robertsonian translocation aberration are the norm, with a population frequency of ∼1:800^16,19,20^. The reproductive consequences for carriers can nonetheless be profound, including increased rates of infertility and aneuploidy or uniparental disomy in their liveborn offspring^11,16^.

Robertsonian translocations are invisible to essentially every diagnostic method other than karyotyping or specialized cytogenetic methods^21^. Sequencing technologies have never been demonstrated to routinely detect these anomalies, and breakpoint positions have been resolved only roughly using FISH and PCR^22^. Ring chromosomes may be suspected via microarray or srGS if there are terminal deletions or duplications^23^, and srGS has enabled breakpoint mapping in a few cases of ring chromosomes subsequent to initial identification by karyotype^24–29^, but neither approach is capable of routine discovery of the diversity of rings that have been observed. Here, we demonstrate that new technologies may offer the opportunity to discover and characterize these anomalies at breakpoint resolution. These advances would resolve breakpoints and rearrangement complexity for individual rearrangements, as well as providing access to novel biological insights into SV formation and functional consequences. Three recent technological developments that could aid in the detection and characterization of SVs in repetitive regions are long-read genome sequencing (lrGS)^30–32^, improved SV discovery algorithms for lrGS^33–35^, and a more complete picture of sequence from the acrocentric p-arms, telomeres, and centromeres as part of the telomere-to-telomere (T2T) reference assembly^17,18,36–39^. In the present work, we test the capacity of lrGS to resolve Robertsonian translocations and ring chromosomes, as well as complex interchromosomal translocations and a complex pericentric inversion. We demonstrate that our analyses were able to discover and resolve these rearrangements, define their sequence complexity, and illuminate mechanisms of formation for these unique classes of SVs.

## Results

### Acquisition and long-read sequencing of a representative set of genome rearrangements

Thirteen samples identified via karyotype to harbor complex rearrangements were selected (Methods) and sequenced with Oxford Nanopore Technologies (ONT) using the R9.4.1 flow cell chemistry to a mean depth of 26x with a median read-length of 9,463 bp (Table 1, Table S1). The dataset included seven ring chromosomes, three Robertsonian translocations, one large inversion, and two inter-chromosomal translocations. Reads were aligned to the T2T chm13 assembly^38^. Rearrangement breakends (Table S2) were detected by a combination of methods: copy number analysis with GATK-gCNV^33^; manual inspection of soft-clipped reads at coverage borders in the integrative genomic viewer (IGV)^40^; SV discovery using pbsv^34^ and Sniffles^35^ lrGS algorithms; and breakend calling via BigClipper, a custom translocation caller that we developed for these studies (Methods).

**Table 1.**
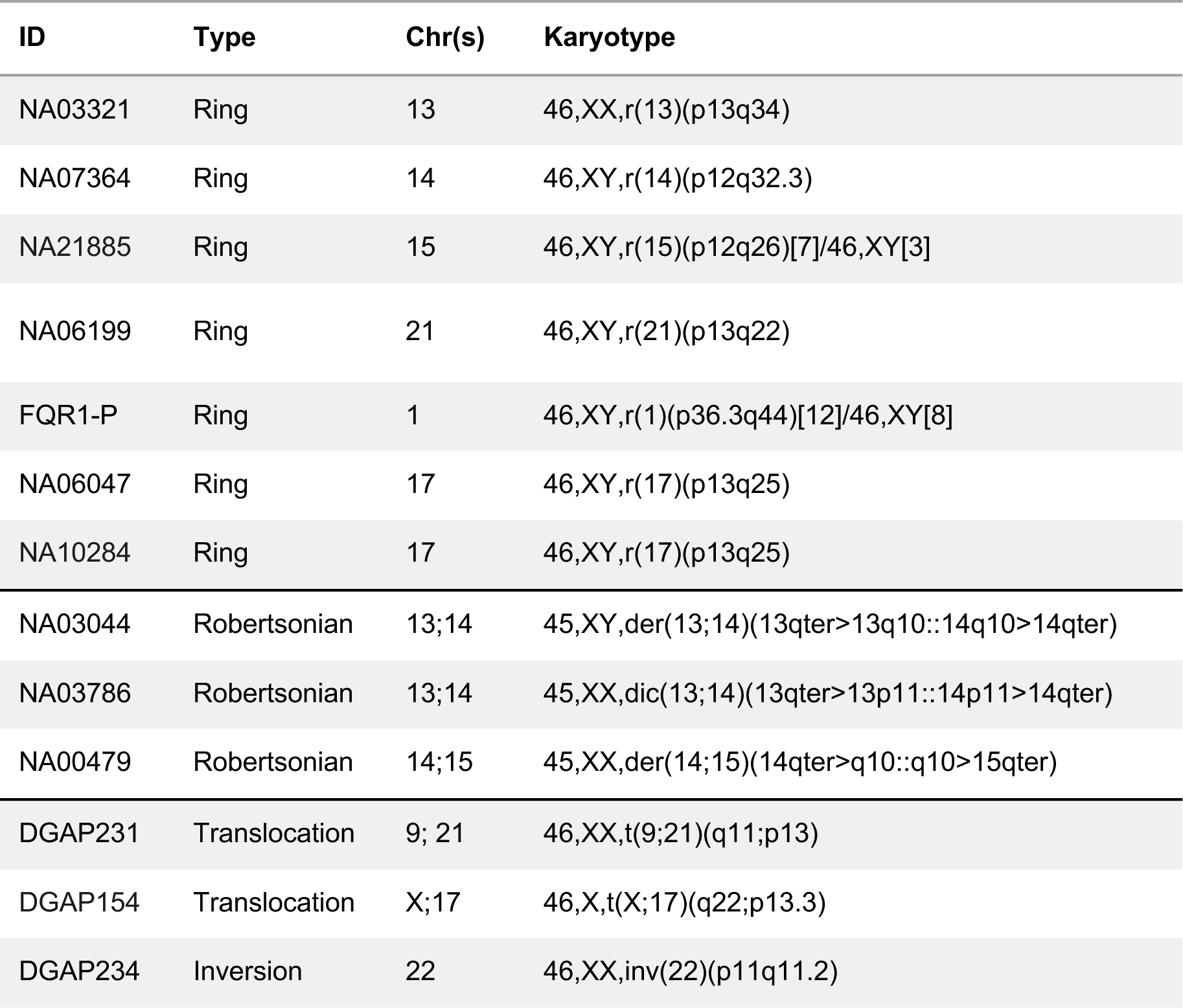
Samples analyzed in this study.

### Ring Chromosomes

Of the seven samples with ring chromosomes, two (NA10284 and NA06047) were rings of chr17. NA10284 is from an individual with intellectual disability (ID), seizures, behavioral disorder, gait abnormality, dysarthria, and short stature. This sample had no cytogenetically detectable change in copy number, but on sequence analysis did exhibit lower coverage in the telomeric 47 kb (T2T chm13 v2.0, RefSeq accession GCF_009914755.1; all subsequent coordinates and intervals use this genome build) of 17p (Fig. 1a). Alignment revealed a group of five clipped reads, whose clipped regions contained copies of the telomere repeat unit (TTAGGG)n, as well as repeats of (TTAAAA)n and (CCAGGG)n, which are known errors made by the ONT basecalling models in telomeric regions^41^. We used the tuned basecaller model from Tan et al.^41^ to re-basecall the clipped reads, resulting in the expected stretches of (TTAGGG)n in each read (Fig. S1a). One read spanned the telomere repeats on both ends, containing 1.1 kp of sequence aligning to chr17:47-48 kb, followed by 4.7 kb of telomere repeats, and ending with 3.7 kb of sequence that aligned via BLAST^42^ to segmentally duplicated subtelomeric regions that included the subtelomere of 17q (chr17:84.270-84.274 Mb) (see Table S2 for RefSeq contig accessions and full breakpoint coordinates). This evidence supported a derived ring structure with a 47 kb p-terminal deletion fused to the end of the full-length q-arm (Fig. 1a).

**Figure 1.**
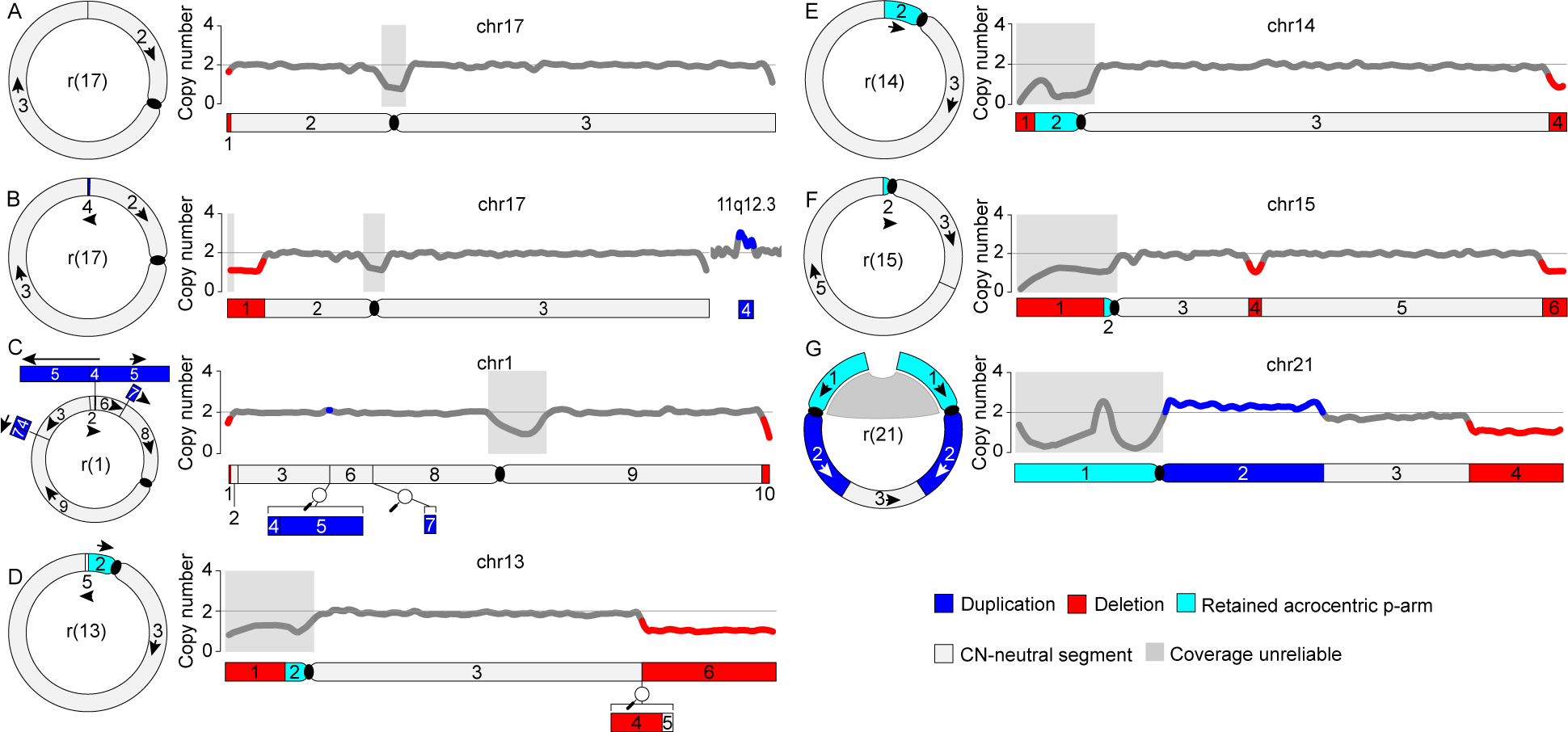
Structures of ring chromosomes. A) Mostly balanced r(17) (NA10284); B) r(17) with terminal p-arm deletion and breakpoint insertion of duplicated sequence from 11q12.3 (NA06047); C) r(1) with terminal deletions as well as interstitial inversions and dispersed duplications. D) r(13) with terminal q-arm deletions and breakpoint in the acrocentric p-arm; E) r(14) with terminal q-arm deletion and breakpoint in the acrocentric p-arm; F) r(15) with terminal and interstitial q-arm deletion and loss of most of the acrocentric p-arm; G) r(21) with terminal q-arm deletion, inversion, and interstitial duplication. Left in each panel, derived structure of the ring chromosome, with segment labels corresponding to the reference chromosome on the right. In panel G, the gray connector between copies of segment 1 indicates uncertainty about their fusion breakpoints. Right in each panel, loess-smoothed estimated copy number of the corresponding reference chromosome. Below, segments relevant to each rearrangement are labeled. Red, deleted segment; blue, duplicated segment; cyan, retained acrocentric p-arm segment with unreliable copy number estimates due to repeat content; gray, segment with no change in copy number. Centromeres are shown as black ovals.

The second r(17) sample (NA06047; Fig. 1b) was collected from an individual with Miller-Dieker syndrome, which is characterized by lissencephaly and characteristic facies and is caused by combined loss of *YWHAE* at chr17:1.23-1.29 Mb and *PAFAH1B1 (LIS1*) at chr17:2.48-2.57 Mb^43,44^. Normalized copy number was estimated with GATK-gCNV and indicated a heterozygous deletion of 17p at chr17:pter-5.76 Mb (Fig. 1b), fully spanning the Miller-Dieker syndrome critical region. At the 1n-2n copy number boundary, a pileup of clipped reads supported a breakend between chr17:5.76 Mb (+) and chr11:62.51 Mb (-) (Fig. S1b). The chr11 breakend marked one end of a 327 kb region with elevated copy number, the other end of which contained a pileup of five clipped reads at chr11:62.84 Mb. The clipped reads at this site contained both canonical and artifactual telomere repeats that converted to canonical telomeres after re-basecalling with the telomere-tuned model (Fig. S1b)^41^. Two reads flanked the telomere repeats on both ends, with one end of each anchored at chr11:62.84 Mb and the other end aligning to subtelomeric sequences including 17q. These data support a derived ring structure with a 5.76 Mb 17p-terminal deletion, insertion of 327 kb of duplicated chr11 material, and fusion to the full-length end of 17q (Fig. 1b). This compares to the karyotype, in which ring 17 was unable to be resolved in this level of detail (Table 1).

The r(1) sample (FQR1-P; Fig. 1c) was collected from an individual with undescended testes, ureter anomaly, ID, progressive contractures of the second digit, small fingers that hyperextend at rest, nail clubbing, short stature, microcephaly, and childhood acute lymphoblastic leukemia in remission following chemotherapy. Karyotyping showed 8/20 normal cells with 12/20 containing the ring (Table 1). The copy number profile of chr1 generated by GATK-gCNV revealed heterozygous terminal deletions as well as an interstitial duplication. Examination of copy number thresholds and clipped read pile-ups identified by BigClipper yielded a complex structure with a total of eight distinct breakpoints, including two inversions and three dispersed duplications (Fig. 1c; Fig. S1c). Sequencing of the proband’s transmitting parent did not reveal any structural anomalies on chr1.

The remaining four rings were of acrocentric chromosomes 13, 14, 15, and 21. The r(13), r(14), and r(15) samples were all found to have terminal deletions on both chromosomal arms. Heterozygous terminal q-arm deletions were apparent in plots of GATK-gCNV normalized copy number (Fig. 1d-g), with a pileup of split reads at each copy number threshold. The r(13) sample (NA03321) was derived from an individual with bilateral cleft lip and palate, microcephaly, microphthalmos, and hypertelorism – shared features with r(13) and 13q deletion syndrome patients^45^. In this sample, the clipped portion of the split reads at the 13q terminal deletion breakpoint contained a 139 bp inversion originating from 626 bp downstream of the breakpoint, followed by sequence aligning to a segmental duplication that is shared between 13p, 14p, and 21p with 99.2-99.4% sequence identity, which is substantially higher than the accuracy of the ONT reads used here (∼92%) (Fig. S1d). The clipped reads aligned to a homologous position on all three copies of the segmental duplication, with three reads aligning to 13p, four reads aligning to 21p, and one read aligning to 14p. The most likely derived structure involves a translocation to 13p following the short deletion and inversion (Fig. 1d).

Both the r(14) and r(15) samples involved a simple translocation embedded in the split reads at their respective copy number thresholds (Fig. S1e,f). The r(14) sample (NA07364) is from an individual with intellectual disability, seizures, limited speech, micrognathia, scoliosis, absent kidney, rib fusion, hypoplastic scrotum, camptodactyly/brachydactyly, and small/thin build. At the 14q deletion breakpoint, this sample contained 11 split reads whose clipped portions aligned to 11 different positions in the rDNA arrays of acrocentric p-arms: four on 14p, three on 13p, and three on 15p, as well as one that aligned to 14p for 7kb and then aligned to 22p. All aligned in the (-) orientation. The most parsimonious derived structure involves a translocation to the rDNA array on 14p followed by an additional inversion, likely elsewhere in the 14p arm, which is as yet undetected (Fig. 1e).

The r(15) sample (NA21885) was derived from an individual with developmental delay, microcephaly, triangular facies and micrognathia, history of dislocated hip repair and leg-length discrepancy, club feet, cryptorchidism, short stature, and delayed bone age. This sample had 12 split reads at the q-arm copy number threshold whose clipped portions aligned within a 104 kb region in a centromeric satellite of 15p (Fig. S1f), suggesting a derived structure involving the loss of most of the 15p arm (Fig. 1f). This sample additionally had an interstitial 2.35 Mb deletion in the q-arm with split reads on either side indicating a simple self-contained deletion (Fig. S1f). This deletion had no apparent connection to the ring structure and, with the available data, it was unclear whether it occurred on the ring or the linear chromosome, with the former being illustrated in Fig. 1f.

The r(21) sample (NA06199) was from an individual with bilateral inguinal hernias and mild developmental delays at 14 months. Sequence analysis showed a triphasic copy number profile on 21q, with an elevated copy number (median=2.3) for 13.2 Mb beginning at the centromere, followed by 9 Mb of intermediate copy number (median=1.8) and 11.6 Mb of low copy number (median=1.0) (Fig. 1g). The departure of the former two values from round copy numbers of 3n and 2n suggests mosaic complexity, likely involving a combination of mosaic ring loss and structural variability in the ring^46^. Split reads at the 2.3n-1.8n threshold had clipped portions that aligned to the 1.8n-1n threshold in inverted orientation (Fig. S1g), consistent with a derived structure where the terminal q-arm breakpoint between segments 3 and 4 attached to an inverted duplication at the end of segment 2. Karyotyping predominantly showed a dicentric ring, so completion of the ring would involve a fusion between the acrocentric p-arms (Fig. 1g), which thus far has not been detected. This derived structure is consistent with previous analyses of the sample^47^.

### Linear Rearrangements: Inversion and Translocations

A pericentric inversion of chromosome 22 (DGAP234) was from an individual with familial IgG3 subclass deficiency with poorly defined inheritance. This sample had no large copy number changes detectable by GATK-gCNV. To further probe this initially apparently balanced rearrangement, we generated SV calls from pbsv^34^, Sniffles^35^, and BigClipper (see Methods), which in some cases was able to detect breakpoints missed by the other callers. We detected two pileups of clipped reads, involving two distinct sets of molecules, within 81 bp of one another on 22q12.2 (segment 8; Fig. 2a). One pileup aligned to the rDNA array present on all acrocentric p-arms (-), representing the distal end of the inversion, while the proximal end on 22p included two additional breakpoints that were resolved by individual reads that spanned consecutive breakpoints, leading to the derived inv(22) structure shown in Fig. 2a.

**Figure 2.**
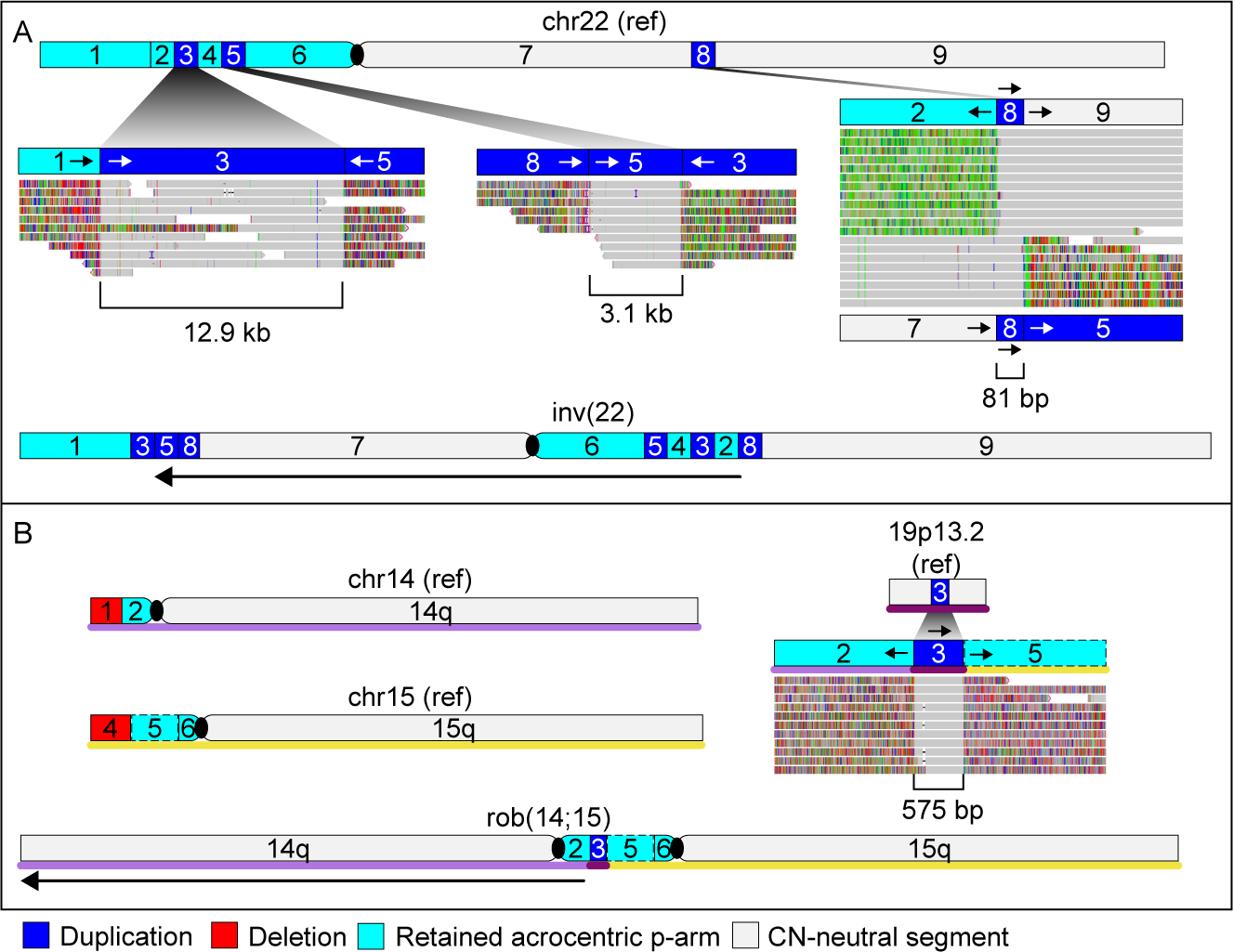
Structures of linear rearrangements. A) Complex pericentric inversion of chromosome 22 (DGAP234). Top, reference chromosome labeled with segments involved in the rearrangement, with cutouts showing IGV screenshots of soft-clipped molecules supporting each breakpoint. Bottom, derived structure of the inverted chromosome. B) Robertsonian translocation between chromosomes 14 and 15 (NA00479). Top, reference chromosomes labeled with segments involved in the rearrangement, with an IGV screenshot centered on segment 3 showing soft-clipped molecules supporting all three chromosomes involved in the translocation. Bottom, derived structure of the translocation. Colored lines below the chromosomes indicate the source chromosome: light purple, chr14; yellow, chr15; dark purple, chr19. The dotted line around segment 5 indicates uncertainty regarding the location of its upstream breakpoint, which occurs in a satellite array spanning ∼8 Mb. Red, deleted segment; blue, duplicated segment; cyan, retained acrocentric p-arm segment with unreliable copy number estimates due to repeat content; gray, segment with no change in copy number. Centromeres are shown as black ovals. Segment sizes are not to scale.

The dataset also contained three Robertsonian translocations, i.e. fusions between acrocentric p-arms (see Table S1 for phenotypes). Copy number ascertainment in acrocentric p-arms was highly variable and generally unreliable, consistent with repeat-associated misalignment as well as known extreme diversity of acrocentric p-arms^48,49^. No breakpoints were identified for the two rob(13;14) samples, but for the rob(14;15) sample (NA00479), BigClipper identified a breakpoint that was unique in the dataset between chr14:5Mb and chr19:10.6Mb (Fig. 2b). This breakpoint was marked by 11 clipped reads, each containing three total alignments: first to 14p, followed by a dispersed duplication of 575 bp from chr19, followed by sequence aligning to different positions on 15p ranging from 7.2 Mb to 13.99 Mb (Fig. 2b, Table S2). The 575 bp segment from chr19 was flanked on both ends by Alu elements (Table S2). The alignments on 15p were all localized within a satellite of class HSat3 subfamily A5 that spans nearly 8 Mb and is the only representative of its subfamily in the T2T assembly. Thus the breakpoint within 15p can be confidently assigned to this satellite, although its exact position cannot be refined further with current data.

Our dataset contained two additional non-Robertsonian translocations, t(9;21) (DGAP 231; phenotype of learning difficulties, anxiety and obsessive compulsive traits, and possible intestinal malformation or malrotation) and t(X;17) (DGAP 154; phenotype of motor and speech delays, facial dysmorphism and possible seizures and obsessive compulsive behaviors). The t(9;21) karyotype described a translocation between the 21p acrocentric arm and the centromere of chr9; no breakpoints were detected for this sample. The t(X;17) sample was analyzed previously with srGS^5,7^ and multiple breakpoints were detected in an apparent chromoanasynthesis event, although missing breakpoints prevented a full reconstruction of the derived chromosomes. Using ONT data, BigClipper detected all of the previously identified breakpoints, revising the position for two breakpoints, and further identified an additional breakpoint junction that was cryptic to the srGS data, potentially due to the AluSx elements present at both breakpoints (new segment boundaries at 22-23 and 19-20; Fig. 3a, Fig. S1h, Table S2). Manual examination of the breakpoint sequences revealed three additional small dispersed duplications (35-144 bp; segments 14, 16, and 18) that were not captured in minimap2 alignments and instead were represented as unaligned portions of reads; these were recovered by using BLAST^42^ to align the insertion sequences to the T2T chm13 reference (Table S2, Fig. S2). The larger set of breakpoints permitted a reconstruction of the derived structure, with two equally parsimonious options (Fig. 3b, Fig. S3). Duplication of segment 5 generates extra copies of *XIAP* and *STAG2*, the critical region for Xq25 duplication syndrome^50,51^, potentially explaining the patient’s delays and dysmorphism.

**Figure 3.**
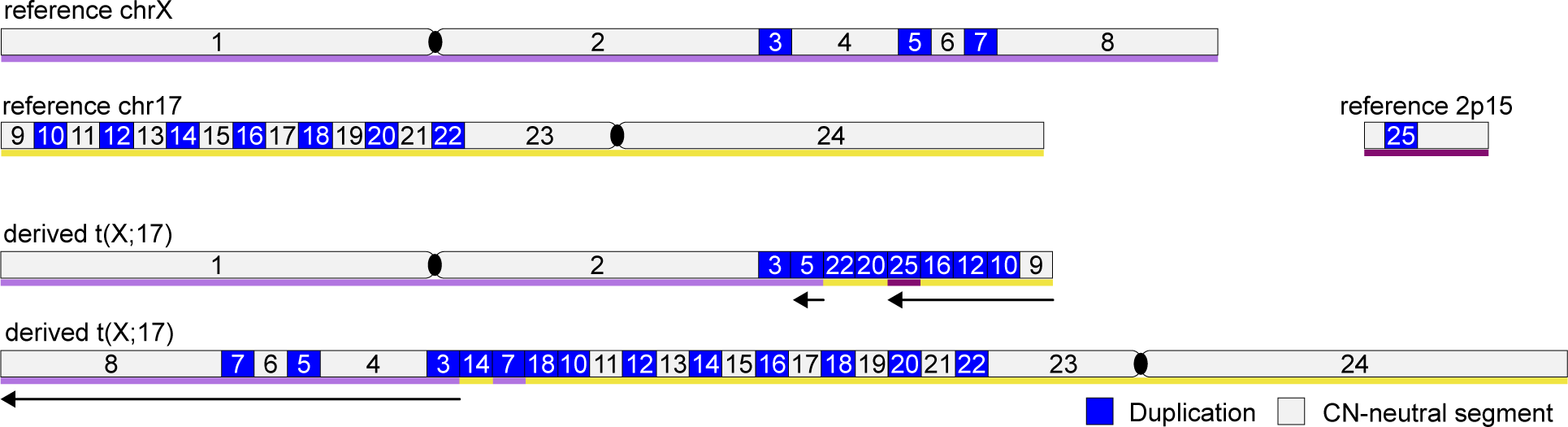
Structure of a translocation with chromoanasynthesis between chrX and chr17 with an inserted duplication from chr2 (DGAP154). Top, reference chromosomes labeled with segments involved in the rearrangement. Bottom, derived structures of the translocated chromosomes. Small segments are not to scale; see Table S2 for breakpoint coordinates. Colored lines below the chromosomes indicate the source chromosome: light purple, chrX; yellow, chr17; dark purple, chr2. Blue, duplicated segment; gray, segment with no change in copy number. Centromeres are shown as black ovals. Segment sizes are not to scale.

### DNA methylation at rearrangement breakpoints

To test whether large-scale genomic rearrangements affect local DNA methylation levels, we used the ONT data to call 5mC methylation at CpG sites for regions extending 1Mb from each breakpoint into the retained segments. Read-based phasing information was used to detect allele-specific methylation for each sample, and regions with recurrent allele-specific methylation were filtered out (see Methods for details). We detected substantial differential methylation near the breakpoints of four samples: DGAP154 (t(X;17)), NA21885 (r(15)), NA10284 (r(17)), and NA06047 (r(17)) (Table S4). The strongest signal was seen in the r(15) sample (Fig. 4), where the most proximal gene upstream of the q-arm ring breakpoint, *ARRDC4*, harbored a CpG island over its promoter that was highly methylated among reads from the ring haplotype (0.84 mean methylated fraction) and was hypomethylated among reads from the unaffected haplotype (0.05 mean methylated fraction, *p* < 1e-20 by Wald test), as well as among all other samples in the dataset (0.02 mean methylated fraction) (Fig. 4C). The other cases involved allele-specific methylation of chr17 in the p-arm (NA10284) and q-arm (NA06047) and of chrX in segment 4 of the t(X;17) sample (Table S4).

**Figure 4.**
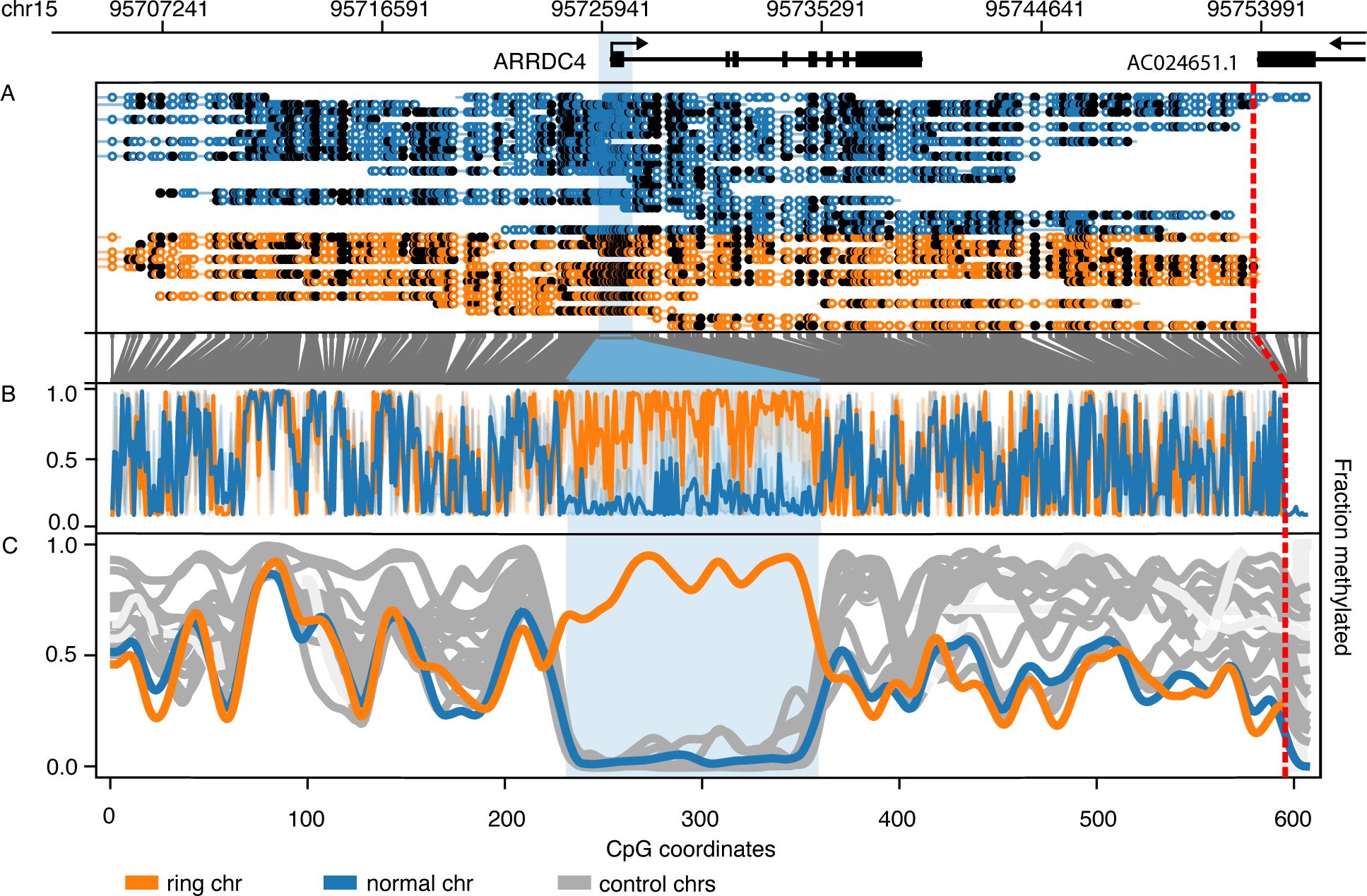
Allele-specific methylation proximal to the r(15) breakpoint of NA21885. From top, gene track showing exons and direction of transcription. A) Phased reads with methylated CpG sites marked with black circles and unmethylated CpG sites marked with open circles, where blue reads are from the unaffected haplotype and orange reads are from the ring haplotype. Below, grey vertical lines mark the conversion from ‘coordinate space’ to ‘CpG space’, in which every position is a CpG site. B) Orange and blue lines show the proportion of reads that were methylated at each site for the ring and unaffected haplotypes, respectively. C) Smoothed depiction of the raw information from B, in addition to gray lines showing the smoothed methylation fraction data from phased unaffected samples (see Methods for details). The transparent blue rectangle highlights the differentially methylated region overlapping the transcription start site of *ARRDC4*. At right, the red dotted line marks the ring chromosome breakpoint.

## Discussion

The advent of long-read sequencing has opened access to sequence-resolved breakpoints for classes of genomic rearrangements that were previously only visible to low-resolution cytogenetic analysis. In this study, we compiled a representative set of 13 rearrangements and used lrGS to characterize their breakpoints. We resolved breakpoints for 10 of 13 samples, including all seven ring chromosomes, one Robersonian translocation, and two of three complex SVs. This work presents, to our knowledge, the first demonstration of a sequence-resolved Robertsonian translocation with lrGS technologies as well as the largest collection of sequence-resolved ring chromosomes to date.

The ring chromosomes evinced a wide range of structural variability. The simplest, r(17) in Fig. 1a, was a nearly complete ring with only one ∼40 kb 17p-terminal deletion fused to telomeres and then subtelomeric sequence aligning to 17q. The other r(17) (Fig. 1b) had an overall similar structure with a larger terminal deletion, but between the p-arm breakpoint and the q-arm-associated telomeres was a ∼300kb dispersed duplication from chr11. Four of the remaining ring chromosomes had terminal deletions at both ends of the chromosome (Fig. 1c-f), with one having additional breakpoint complexity in the form of a small deletion and inversion (Fig. 1d). The ring of chr1 was highly complex at a scale that was cryptic to karyotyping, having a total of eight distinct breakpoints, three dispersed duplications, and two inversions (Fig. 1c). The r(21) sample, in addition to harboring a large inverted duplication, displayed mosaic complexity, directly observed with karyotyping^46^ and reflected in the long-read data with copy numbers that departed substantially from the expected round values (Fig. 1g). The derived structure in Fig. 1g was consistent with the predominant form of the ring seen in karyotypes^46^ and refined with molecular cloning^47^.

Nine of the samples in our dataset had rearrangements involving p-arms of acrocentric chromosomes. While these regions now have a haploid reference assembly^38^, their highly repetitive content continues to present challenges in long-read alignment and breakpoint mapping. The most challenging rearrangements were those where both ends of a breakpoint were located in an acrocentric p-arm or similarly long repetitive region. Of the three Robertsonian translocations, we were unable to determine breakpoints for two of them, while the third had a breakpoint with a translocation from a third chromosome (Fig. 2b). The two unsolved Robertsonian translocations were of karyotype rob(13;14), which have been proposed to fuse inside of the multi-Mb >99% sequence identity segmental duplication shared between 13p, 14p, and 21p^52^. Another challenging unsolved case involved a translocation between 21p and the pericentromeric region of chr9, which contains a 28 Mb satellite array^36^. Furthermore, the r(21) sample had resolved breakpoints in 21q, but breakpoints from a putative 21p-21p fusion remained elusive (Fig. 1g). In contrast, we were able to characterize acrocentric 22p internal rearrangements involved in the pericentric inversion of chr22 (Fig. 2a) because single long-reads spanned multiple segments including a region of 22q (Fig. S1).

Furthermore, the five p-arms of acrocentric chromosomes have high similarity – and interchangeability owing to inter-chromosomal exchange between non-homologous acrocentric p-arms^52,53^ – of sequence that can confound precise definition of the originating chromosome for a given read. Of the three acrocentric ring chromosomes with resolved p-arm breakpoints, two had breakpoint-associated clipped reads aligned to multiple chromosomes. The r(13) breakpoint was located in a segmental duplication shared between 13p, 14p, and 21p, while the r(14) breakpoint was inside of the rDNA array that is shared between all five acrocentric p-arms. The inv(22) sample also had breakpoint-associated clipped reads aligning to the rDNA array. In contrast, the r(15) breakpoint clipped reads aligned to multiple positions that were all located in a 15p centromeric satellite, restricting it to a single chromosome. Similarly, in the rob(14;15) sample, breakpoint-associated clipped reads aligned, on one end, exclusively to one position on 14p, and on the other end within an 8Mb satellite of a subclass present only on chr15.

Our results reveal considerable heterogeneity and new insights into breakpoint localization and rearrangement mechanisms. For example, the ring chromosome breakpoints often involved additional complexity, reminiscent of our earlier sequencing studies of balanced chromosomal translocations and inversions^5,7,54^. 39% (12/31) of breakpoint junctions had 3-6 bp of microhomology (Fig. S2, Table S3), consistent with microhomology-mediated repair mechanisms such as microhomology-mediated break-induced replication (MMBIR)^55,56^ or microhomology-mediated end-joining (MMEJ)^57^. Other breakpoints had no observed microhomology (Fig. S2, Table S3) and were more consistent with a non-homologous end joining (NHEJ) mechanism^58^. The t(X;17) chromoanasynthesis case had *Alu* elements overlapping 11 of 22 breakpoints (Table S2), some with long stretches of homology (Table S3), suggesting the partial involvement of non-allelic *Alu/Alu* homologous recombination^59^ in this case. The resolved Robertsonian translocation (rob(14;15)) had breakpoints consistent with a microhomology-mediated mechanism, although the more common rob(13;14) and rob(14;21) rearrangements are likely caused by NAHR between long segmental duplications^52^. While we assessed an intentionally diverse set of rearrangements, future application of our methods to larger sets of specific rearrangement types (and with T2T references beyond a single haploid assembly) is poised to contribute answers to additional long-standing questions about their formation and biology, for example: the preference for certain chromosomes in Robertsonian translocations^52,53^; the properties of fused or dicentric centromeres^52,60^; the frequency of fusion genes/enhancer swapping; position effects on gene expression^61–64^; and the biology of subsequent mitotic/meiotic errors (i.e. “tissue-specific dynamic mosaicism”)^65^.

We used the ONT data to detect 5mC CpG methylation in regions around rearrangement breakpoints, and identified unique allele-specific DNA methylation in four rearrangements: r(15), both r(17)s, and t(X;17). In r(15), the strong hypermethylation signal on the q-arm of the ring haplotype proximal to the breakpoint was likely related to the location of the p-arm breakpoint, which was inside of an Alpha satellite in close proximity to the start of the centromere - a region known to be heavily methylated^66^. Bringing this pericentromeric region in close proximity to the q-arm may have caused the spreading of heterochromatin to the q-arm. In contrast, the ring samples in our dataset that did not have substantial allele-specific methylation also had p-arm breakpoints much farther from the centromere. Interestingly, both r(17) samples showed signs of allele-specific methylation, one near the p-arm breakpoint and the other near the q-arm breakpoint. These two ring samples were unique in our dataset for retaining telomeres at their breakpoints, which have previously been associated with regional down-regulation of gene expression in cases of r(17)^64^. The final sample with allele-specific methylation had the t(X;17) rearrangement and was differentially methylated at five breakpoint-proximal regions on the X chromosome. While X chromosome methylation signal may be difficult to disambiguate from the process of X inactivation (Xi), our finding suggests some disruption of DNA methylation on chrX associated with the translocation, including *STAG2* hypermethylation that could potentially abrogate the effects of Xq25 duplication.

The ability to detect and resolve repeat-localized structural variants using a sequencing methodology is of obvious potential clinical utility. The specific gene content in the case of deletions/duplications and structure in the case of gene breakage events can be essential to explaining or predicting phenotypes resulting from structural variation. Furthermore, the detection of rearrangements with balanced genic content (e.g. Robertsonian translocations, complete ring chromosomes) is of high importance given the significant potential reproductive risk in asymptomatic carriers and inability to be identified by the most common genetic screening methods (microarray and exome sequencing). Finally, the clinical utility of methylation data has recently become apparent^67^; to ascertain methylation status genome-wide and layer this on a structural variant map holds considerable clinical potential^68^.

There are limitations to these studies that could be improved upon in future implementations. For example, we focused on a single long-read method, ONT, while others remain to be tested such as optical mapping and PacBio HiFi sequencing^60^. This point is reinforced by earlier assembly-based methods for SV discovery and the T2T assembly itself^38^, all of which required multiple complementary technologies and analytic methods. Nonetheless, these data suggest that routine sequencing and discovery of these rearrangements is tractable when a karyotype is accessible. The rarity of ring chromosomes provides a considerable challenge to perform benchmarking of the likelihood of *de novo* discovery, though future studies could pursue blinded benchmarking.

In summary, we demonstrate that long-read sequencing and mapping to the T2T reference enables breakpoint localization of historically difficult-to-resolve genome rearrangements including ring chromosomes, a Robertsonian translocation, a complex interchromosomal translocation, and a complex pericentromeric inversion. Future work will ideally test additional and improved long-read methods and analyses and additional samples to arrive at robust, blinded detection of these rearrangements. Our work progresses toward the goal of a single genetic test to identify all genetic variation.

## Methods

### Samples

Table S1 details the samples used in this study, including their origin, known cytogenetic diagnoses via karyotype (+/-microarray, FISH), quality control data, and subject phenotypes. Samples were selected to be representative of various ring chromosomes, Robertsonian translocations, and other rearrangements of interest, favoring samples of highest DNA quality. Samples from the Coriell Institute for Medical Research (https://www.coriell.org/) were used with permission of the Coriell Repository Review Board. Samples from the Developmental Genome Anatomy Project (DGAP; Higgins AJHG 2008) were enrolled under Partners HealthCare IRB protocol 1999P003090. Samples from UCLA, including probands and available parents, were enrolled under IRB protocol 11-001087.

DNA from Coriell was prepared from fibroblast or lymphoblastoid cell lines, as described here: https://www.coriell.org/0/Sections/Support/Global/DNA.aspx?PgId=689 and https://www.coriell.org/0/Sections/Support/Global/QCdna.aspx?PgId=410. The DNA was prepared with standard methods to replicate current clinical diagnostic lab practices rather than preparations tuned for ultra-long reads^69^. Briefly, DNA was isolated using an automated magnetic bead purification method (Hamilton Microlab STAR liquid handling machine and Promega ReliaPrep Large Volume HT gDNA isolation system) or manually via the modified Miller’s salting out procedure. DNA from UCLA was prepared from whole blood via the DNeasy Blood and Tissue Kit (Qiagen, Hilden, Germany). DNA for DGAP samples was obtained from blood or lymphoblastoid cell lines. DNA quality was assessed via the Tapestation 2200 or 4200 machine (Agilent Technologies, Santa Clara, CA) and Tapestation Analysis Software A.02.02 or 3.2.

### Sample processing

Long-read sequencing was conducted under MassGeneralBrigham IRB 2013P000323. For ONT library preparation, ≥2 ug of high molecular weight genomic DNA (more than 50% of fragments ≥40 kb) was sheared to ∼20 kb using the Megaruptor (Diagenode B06010002), followed by DNA repair and ligation of ONT adapters using the ONT Ligation Sequencing Kit (SQK-LSK109). Short fragments were removed in a size selective bead cleanup included in the kit. The ligated libraries were quantified using the Qubit dsDNA High Sensitivity assay (Thermo Q32854) and libraries were prepared for sequencing using ONT provided buffers (EXP-AUX002). Sequencing was performed on the PromethION instrument using R9.4.1 flowcells (FLO-PRO002) and utilized ONT’s on board high accuracy basecalling in real time.

Basecalled reads were processed with Broad’s long reads pipeline in the Terra cloud-based platform (https://app.terra.bio/). Reads were aligned to the T2T chm13 assembly^38^ (initially v1.0, then v1.1 and v2.0 as they became available) using minimap2^70^ v2.17. SVs were discovered using Sniffles^35^ v1.0.12 and pbsv^34^ v2.6.0.

### Breakpoint discovery

To search for breakpoints, we first used R^71^ to visualize the copy number profile of the relevant chromosome for each sample as determined by GATK-gCNV^33^. The borders of apparent copy number changes were examined in IGV^40^ for pile-ups of clipped reads. Clipped reads were further visualized with Ribbon^72^. Breakpoint coordinates are genome build T2T v2.0^38^ unless otherwise stated. Several of the ring chromosome samples needed further analysis to identify and refine their breakpoints. For example, each r(17) sample had clipped reads where the clipped portion resembled telomeric repeats. We re-processed clipped reads resembling telomeric repeats with an ONT basecaller tuned to fix a telomere-specific base-calling issue^41^ in the Nanopore basecalling models and confirmed that these reads contained telomere sequence. Then, alignment to the T2T assembly using BLAST^42^ further localized the clipped sequence to subtelomeric segmental duplications present at the end of 17q as well as other subtelomeres.

To augment SV discovery from Sniffles and pbsv for relatively balanced translocations, we wrote a custom breakpoint-finder called BigClipper. Briefly, the method read the BAM files and extracted reads with supplementary alignments that had at least 500 bp of soft-clipping. Retained reads within 5 bp of one another were linked into clusters. Clusters with at least five (by default) reads were retained, and their supplementary alignments were then clustered by position, grouping those that were within 50 bp. Clusters with more than ten (by default) supplementary alignment groups were discarded as false positives. The remaining clusters were output in VCF format and manually analyzed in the regions of interest. For cases where breakpoints could not be found with default settings, we lowered the minimum number of reads per cluster from 5 to 2 and increased the maximum number of supplementary read clusters from 10 to 20. Using BigClipper, we were able to identify clipped-read-based breakpoints in r(1), inv(22), rob(14;15), and t(X;17) (Figs 1-3). BigClipper source code is available at https://github.com/yuliamostovoy/bigclipper.

### Breakpoint sequence analysis

To characterize homology and additional complexity at breakpoint junctions, sequences flanking the reference breakpoints were extracted using seqtk^73^. To obtain the derived breakpoint junction sequences, breakpoint-spanning reads were extracted from BAM files, converted to fasta format with SAMtools^74^, and assembled using Flye^75^. Contigs were then aligned to the reference using minimap2^70^, and sequences flanking the breakpoints were extracted using seqtk. In some cases, assembly of the breakpoint-spanning reads failed, in which case raw reads were used as the source for the junction sequence. Derived and reference sequences around the breakpoints were manually aligned to identify homology or insertions between the breakpoints (Fig. S2, Table S3). Inserted sequences were aligned with BLAST^76^ to the T2T chm13 reference and the Dfam^77^ consensus sequences of human repeat elements.

### Methylation

To detect methylation in the vicinity of rearrangement breakpoints, we extracted fast5 data for reads that mapped to a window extending 1 Mb into each retained segment. For the selected reads, 5mC methylation was called at CpG sites using Megalodon with the “res_dna_r941_prom_modbases_5mC_CpG_v001.cfg” basecalling model. Per-read modified base results were output in database form with the “--outputs per_read_variants” option, converted to text form using “megalodon_extras per_read_text modified_bases,” and finally converted to a form parseable by Methylartist^78^ using the “methylartist db-megalodon” command. Separately, haplotagged BAM files were produced using the PEPPER-Margin-DeepVariant pipeline^79^. Per-read modified base information and haplotagged BAMs were input to Methylartist in order to produce phased visualization (with the “locus” command) as well as text files (using “wgmeth --dss”, with the “--motif CG --phased -m m” flags used in both cases) that were input to DSS to detect differentially methylated regions^80^. Using the DSS library in R, differentially methylated regions (DMRs) were called using the following command: callDMR(dmlTest, minCG = 5, p.threshold=1e-20, delta=0.2). These DMRs were exported to a BED file and bedtools^81^ was used to filter DMRs that showed allele-specific differential methylation across multiple samples, i.e. to remove recurrent imprinted regions. The exception to the uniqueness filter was that samples with the same karyotype were allowed to share DMRs (i.e. the two r(17) samples and the two rob(13;14) samples). The final set of DMRs was filtered to exclude regions with fewer than five phased supporting reads.

## Acknowledgements

These studies and personnel were supported by grants from the National Institutes of Health: HD081256, HD096326, HD104224, MH115957, HG011755, and K08NS117891, and the Simons Foundation for Autism Research Initiative (SFARI #573206). The authors thank Serkan Erdin for fruitful advice and discussion.

