## Supplementary figures and images for "Resolution of ring chromosomes, Robertsonian translocations, and complex structural variants from long-read sequencing and telomere-to-telomere assembly"

### Supplemental Figure 1

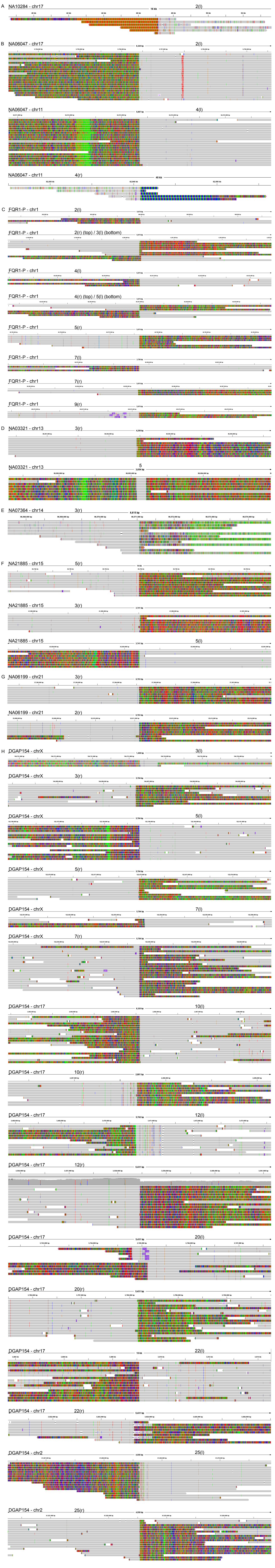

### Supplemental Figure 3

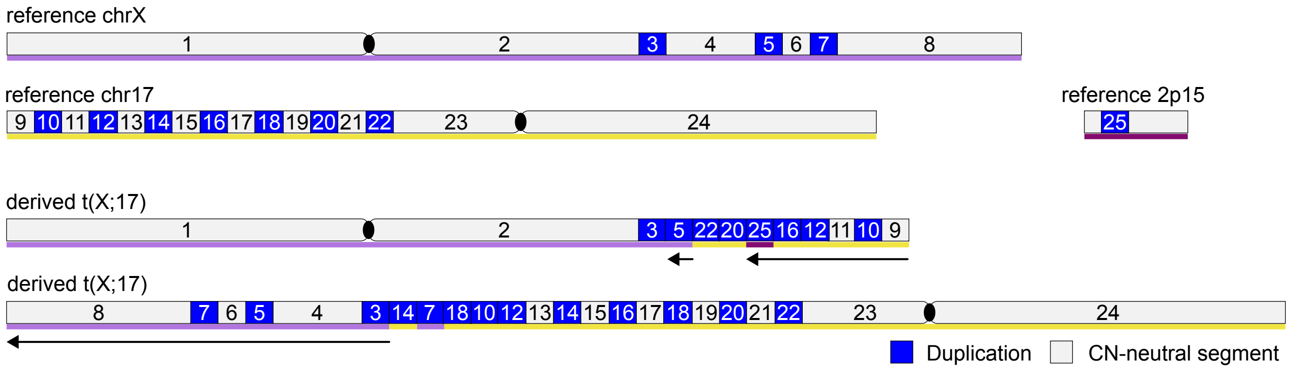
