## Supplemental Figure 2 for "Resolution of ring chromosomes, Robertsonian translocations, and complex structural variants from long-read sequencing and telomere-to-telomere assembly"

```
A  NA10284 r(17)
   read-f5169c48…:8240_8277     – CTTGGCAGGACGGTGAGTGGCC-TTCTCTCCATGCTCCT
   NC_060941.1:g.47082_47120 2l+ CCCATGAGGAGAGAGGAAGGCCTTTCTCTCCATGCTCCT

B  NA06047 r(17)
   contig_1:23773_23820                + GCCAATCCATTGGCTTGGCTGGGCTATCTATGTTGATGTCTCCTTTTG
   NC_060941.1:g.5756391_5756430   2l-          ACTGGCTTGGCTGGGCTATCCAGAGTCTTCCTCCTACAAG
   NC_060935.1:g.62513324_62513363 4l+ GGCCCTCAATGTTCATACTTGGGCCCTCTATGTTGATGTC

   read-33f2c5e2…:15062_15101          + TTAAACCTCTACTAAAATCCTAACCTAACCCCTAACTTGC
   NC_060935.1:g.62840884_62840923 4r+ TGAAACCTCTACTAAAATCCCAATAACTAGGCTTGCTTGG

C  FQR1-P (r(1))
   read-0697ed0c…:1617_1656              + ACTCCCGCAGAATTAGTGTATCCATCTAGGTCCCAGGTCC
   NC_060925.1:g.4200300_4200339      3l- ACTCCCGCAGAATTAGTGTATTATCCATAAAGACTGGGGT
   NC_060925.1:g.843949_843988        2l+ AGGTCCATCTAGGTCCCAGGTCCATCTAGGTCCCAGGTCC

   read-f20275dc…:7926_7964              + TAGTTATTCGGCTTC-AATAACTACATTGAATAGGAGTGG
   NC_060925.1:g.4200264_4200303      2r+ TAGTTATTCGGCTTC-AATAAGTAAATAAAATGATAAACCC
   NC_060925.1:g.46757591_46757628    5l- CCTGGCCAGAACTTCCAAC-ACTACATTGAATAGGAGTG

   read-adbbc046…:7380_7419              + GGCAGGTGCTGTAATGCCAGACCTGTGCACTGCTTCAGAT
   NC_060925.1:g.46436978_46437016    5l- GGCAGGTGCTGTAATGCCAGCTACTTGGGAGGCTGAGGC
   NC_060925.1:g.46430685_46430724    4l+ GGGATGGGGGTGGCAGAGGCACCTGTGCACTGCTTCAGAT

   read-6bc5c4e1…:2611_2650              + TCAGCCTCCCAAGTAGCTGGAGTTGGTGAAAATATCATTT
   NC_060925.1:g.46436981_46437020    4r+ TCAGCCTCCCAAGTAGCTGGCATTACAGCACCTGCCACCA
   NC_060925.1:g.66579846_66579885    7l+  CACCATGCAAGGAGTACCTAGTTGGTGAAAATATCATTTT

   read-233a2cad…:821_859                + TCTGCAGACACATCTGAATAATAAATACATGGATTTTTT
   NC_060925.1:g.66585210_66585249    7r+ TCTGCAGACACATCTGAATCACTTCTGGCCATCAGCAGTT
   NC_060925.1:g.245674130_245674167 9r- TGAGAAAAAAGCTACTTTTAATAAATGCATGGATTTTT

D  NA03321 r(13)
   contig_1:2164_2203                  - TTTTAAGGTGTTTACCCAAATTCTAAAATTATAAATAAGT
   NC_060937.1:g.86092431_86092451 3r+ TTTTAAGGTGTTTACCCAAACCTCTAAGCATAAAATAAAT
   NC_060937.1:g.86093198_86093211 5r- GAGGGAACCATGACCCCAAATTCTAAAATTATA

   contig_1:2030_2069                  - CCTGTTATTTGTGCAACGGTGCGTCTGATCTCCGATGCTT
   NC_060937.1:g.86093078_86093087 5l- CCTGTTATTTGTGCAACGGTAGTTAAATCATTTTTATTG
   NC_060937.1:g.11712683_11712691 2l+ CAAGCCCCATGGAGTGTGGTGCGTCTGATCTCCGATGC
   NC_060945.1:g.8774292_8774297   2l+ CAAGCCCCATGGAGTGTGGTGCGTCTGATCTCCGA

E  NA07364 r(14)
   contig_1:8905_8944                  + CACAGAAGACGGTTTAAAATGAAACAAAACAAGCCAACAA
   NC_060938.1:g.98571043_98571080 3r+ CACAGAAGACGGTTTAAAATGAAAAGAAGCTTCTGGGA
   NC_060938.1:g.2222120_2222158   2l- CGAACAAAACAAAACAGAACGAAACAAAACAAGCCAACA

F  NA21885 r(15)
   contig_1:4950_4989                  - TGAAAGCAGACTAATACAGGCATTCTGAGAAACTTCCTCG
   NC_060939.1:g.95753886_95753924 5r+ TGAAAGCAGACTAATACAGGTACGGTCATGTGTGTGTGT
   NC_060939.1:g.15547796_15547834 2l+ AAATGAAAAGCACACAGAGGCATTCTGAGAAACTTCCT

   contig_1:15752_15810                + CGCCTTGGCCTCCCATAGTGGAGCTAACAATATGAAAAGTGTTCGGTTCTTTTT-GAAAT
   NC_060939.1:g.41589498_41589537 4l+ CGCCTTGGCCTCCCATAGTGCTAGGATTACAAGCGTGAGC
   NC_060939.1:g.43939191_43939230 4r+                     TTTTCAGCTCTATGAATTTTGTTCGGTTCTTTTTTGAAAT

G  NA06199 r(21)
   contig_1:11595_11643                - CAGCTGGAGGTCAGAAAAGGCATTCACCAGCAATTTTGCTTCCCTAACA
   NC_060945.1:g.37394554_37394590 3r+ CAGCTGGAGGTCAGAAAAGGGTGTTTGTTCAGGGCTG
   NC_060945.1:g.25011103_25011142 2r-              AAGATTTGCTTGGCATGCAATTTTGCTTCCCTAACATCTT

H  NA00479 rob(14;15)
   read-6a0b2f0c…:4550_4593            - TTTCTGTGCTTTGCCAACAGAGGGCTCACTGCAGCCTAGACCTC
   NC_060938.1:g.4974892_4974931   2l-    CTAGGCTTTGCCAACAGAGGGTATTGTAACATATCTCTGC
   NC_060943.1:g.10598738_10598777 3l+ TGGAGTGCAGTGGCATGATCAGGGCTCACTGTAGCCTAGA

   read-6a0b2f0c…:3978_4021.           - ACTAAAAATACAAAAATTAGCCAGTCTATTCCACCAATCCATTC
   NC_060943.1:g.10599313_10599352 3r+     AAAATACAAAAATTAGCCAGGCGTGCTGGCGGGGATCTAT
   NC_060939.1:g.6640479_6640518   5l+  CATTCCATACTATTGCATTCCATTCTATTACATTATATTC

I  DGAP234 (inv22)
   contig_1:17631_17670                - CTCTAAGGTAGGCAGGTCAGTTCTCTGTCTGTCTGTCGGT
   NC_060946.1:g.30858522_30858561 8l- CTCTAAGGTAGGCAGGTCAGGTGACTTATTGAGGAGCCTG
   NC_060938.1:g.2222694_2222733   2l+  GTCTTCTGTCTTACTCCCTTTCTCTGTCTGTCTGTCGGTC

   contig_1:26037_26076                + CTTGTGAGCTCTAAGTTGCTCACACAATAAAAATATACAA
   NC_060946.1:g.30858603_30858642 8r+ CTTGTGAGCTCTAAGTTGCTGAACCAATATAAGAGGCTGT
   NC_060946.1:g.6711555_6711594   5l+  CTGTTCCCAATTTCTTGTCCACACAATAAAAAATATACAA

   contig_1:29575_29614                +  ACTCCCCAGTCTGCCTAAGAAAAAAAAAAAAAA-GGACTGTA
   NC_060946.1:g.6715110_6715149   5r+   CTCCCCAGTCTGCCTAAGATATTTTCTGTTATTTTATAAG
   NC_060946.1:g.6710640_6710679   3r- AAAAAAAAAAAAAAAGAAAAGAAAAAAAAAAAAAAAGGACTG

   contig_1:42438_42482                - TCGTGTTTCTTTCCTTCCCGTCTGTTCCTGCATTGGTTTGCTACA
   NC_060938.1:g.2146368_2146407   1r+       TTCTTTCCTTCCCGTCTGTCTTTTAAAAAATGGAGTGTTT
   NC_060946.1:g.6697696_6697735   3l+ AAACGTATGGTATTTAGTTTTCTGTTCCTGCATTGGTTTG

J  DGAP154 (t(X;17))
   contig_1:7002_7051                    + TATGTAATAGTATATATACTGTTAGTTCAGGCTGGCAGGGTCCAGAAGAT
   NC_060947.1:g.104956693_104956732 3r+ TATGTAATAGTATATATACTATATATATACTGTATATATA
   NC_060947.1:g.122471932_122471971 5r-           ACTTTAGACCTGCTCATCCCAGCTGGGAGGGTCCAGAAGA

   contig_1:7695_7744                    + TGCTCCACCATGCCCGCCTAGGACCTGCCTGGCGGACGGATCACAAGGTC
   NC_060941.1:g.3809081_3809120     22l- TGCTCCACCATGCCCGCCTAATTTTTTTTTCTGTATATTT
   NC_060947.1:g.122158751_122158790 5l+             CAGCACTTTGGGAGGCTGAGGCGGACGGATCACAAGGTCA

   read-c7b75cfc…:3411_3501              + ATCACCACGTGCCTGGTTCATTTTTATTTTTATTTGGTAGAGATGGAGTCTCACTCTGTCGCCCAGGCTGG-GTGCAGTGT-GCAGTCTCG-CT
   NC_060941.1:g.3823624_3823703     22r+             CCTGGTTCATTTTTATTTTTATTTGGTAGAGATGGGGTCTCACTGTATTGCCCAGGCTGGTCTCAAACTCCTGGGCTCAA
   NC_060941.1:g.3745100_3745189     20l+ ACCTAAAACTGCTAAAATAATTTTT-TTTTT-TTTTT--GAGATGGAGTCTCACTCTGTCGCCCAGGCTGGAGTGCAGTGGCGCAATCTCGGCT

   read-8c847b7b:4794_4926               + AAAAAAGAGTTGGGGCCAGGAAGATATGGAAAATGCAGAAAGTTATGCTTGCAGAGTATACTCCACTGTATATATACTATATACACTATATATACAGCCCCTATTCATGCGCCAAGAAAGAAAAAGGCTGTTG
   NC_060941.1:g.3788153_3788192     20r+ AAAAAAGAGTTGGGGCCAGGCACGGTGGTCACGCCTGTAA
   NC_060926.1:g.61922519_61922558   25r-                                                                                                 TATACTCCACCTTGTGTCTGAAAGAAAGAAAAAGGCTGTTG

   contig_1:11362_11424                  - TCACTGTAACCTCCACCTCCTGAGTTCAAGTGATTCTCCTGCTTCAGCCTCCCGAGTGGCTGG
   NC_060941.1:g.3651570_3651632     16r+ TCACTGTAACCTCCACCTCCTGGGTTCAAGCGATTCTTCTGCCTCAGCCTCCCAAATAGCTGG
   NC_060926.1:g.61668659_61668721   25l+ TCACTGCAACCTCCGCCTCCCGGGTTCAAGCAGTTCTCCTGCTTCAGCCTCCCGAGTGGCTGG

   contig_1:11513_11609                  - ATCACTTGAACCCAGGAGGCCCTGGCCTCCCGAGTAGCTGGGATTACAGGCATGCGCCACCCTGAAAACCTCACCAGCAGCCTCCCAAAGTGTTGGG
   NC_060941.1:g.3538442_3538481     12r+ ATCACTTGAACCCAGGAGGCGGAGGTTGGGGTGAGCCTAG
   NC_060941.1:g.3651378_3651935     16l+                                                          GACCCATGATCGGCCCACCTCAGCCTCCCAAAGTGTTGGG

   read-97d3db3d…:1062_1706              + AAAAAAAAAAAAAAAAAAAAAAGG-C-GGGCACAGTGGCT…AAAAAAAAAAAAAAAAAAA-------------TCACAGTGGAGCCTGGGCAT
   NC_060941.1:g.2297059_2297712     10r+ AAAAAAAAAAAAAAAAAAAAAAGGGCCGGGCGCAGTGGCT…AAAAAAAAAAAAAAAAAAAGAAAAGAAAAAGGAAAGTGAA
   NC_060941.1:g.3470617_3471280     12l+ GGGGATAAAAATCACAATAGAAGG-CTGGGCACAGTGGCT…AAAAAAAAAAAAAAAAAAAAAAAAAAAAAAAATCACAGTGGAGCCTGGGCAT

   read-0ecdf091…:5457_5530              + ACTGC--ACA--TCTTTTGATGTACCTAAACCTTGGATCCTTGGGTGCAGCGGCAGAATTGCAGTAAGGCGAATG-GCC
   NC_060947.1:g.104721839_104721879 3l- ACTGCCCACACATCTTTTGAAACATTACAGTGTATTTTAA
   NC_060941.1:g.3570306_3570345     14l+                                       GGTTCAGGCCGTAGCTGGAGTGCAGTAAGGCGAATGACCC

   read-0ecdf091…:5539_5579              + CCTGCACC--CTGTCCCAGGCTATCGCCCAGAGCTGG-GTGCAA
   NC_060941.1:g.3570355_3570395     14r+   TGCACCCACTGTCCCAGGCTGTGGGGGTGCTGAGGGAGTA
   NC_060947.1:g.122553606_122553647 7l+   TTGAGACAGAGTCTTGCTCTGTCGCCCAG-GCTGGAGTGCAG

   contig_1:7455_7512                    + AACTGCACATTCAAAGAGTGCCAGGCTGGTCTCGAACTCCTCTAGTTCCTCCAAACCC
   NC_060947.1:g.122637749_122637788 7r+ AACTGCACATTCAAAGAGTGAACAGCAGTTAAAATGAATG
   NC_060941.1:g.3660561_3660600     18l+                    CACCCCTCCCGGCTTCAGACCCTCTAGTTCCTCCAAACCC

   contig_1:7408_7562                    + AACCCTCAGTGCCTGGGTGACAAGTGTTAGCTAGGATATGAGCAGCAAGACACCA
   NC_060941.1:g.3660596_3660636     18r+ AACCCTCAGTGCCTGGGTGACGAGGCACCACCACACTCTCC
   NC_060941.1:g.2065576_2065615     10l+                  CCACCATTCCAGGAGTTAAGATATGAGCAGCAAGACACCA
```
